# Multistage AI Crop Inspection and Segmentation Framework (MACISF)

**DOI:** 10.64898/2026.10.11.758275

**Authors:** Prithul Joshi, Aryaman Nagrath

## Abstract

Crop quality assessment is one of the most critical components of modern agriculture, which directly influences yield optimization, post-harvest management, and food security. Conventional evaluation of field crops relies on manual inspection, which is time-consuming, subjective, and unscalable. This paper proposes the Multistage AI Crop Inspection and Segmentation Framework (MACISF), an automated multistage deep learning pipeline for fruit quality evaluation from visual features. The pipeline is comprised of three stages: (i) an EfficientNetV2-L backbone, fine-tuned with a three-stage progressive strategy, for crop type classification; (ii) seven species-specific EfficientNetV2-S classifiers for healthy-versus-defective detection; and (iii) a YOLOv8m module for defect localization. Trained and evaluated on a composite dataset spanning seven fruit species, MACISF achieves 99.23% crop type accuracy (macro F1-score 99.23%, macro AUC-ROC 0.993), 99.39% mean health state accuracy with 99.51% mean defective-class recall, and a projected grading mAP@0.5 of 92.3%. On crop type classification, the Stage 1 classifier is second only to InceptionV3 (by 0.15 percentage points) among seven models trained on the same data, and outperforms ResNet-50, Xception, EfficientNetB3, DenseNet-121, and MobileNetV2. Unlike these single-task baseline models, MACISF additionally delivers health-state assessment, defect severity grading, and a per-unit economic and nutritional evaluation. Results are reported on held-out test data from a controlled dataset, and field validation is left for future work.

## 1 Introduction

The agricultural production plays a very important role in global food security and economic stability. Agriculture is one of the major contributors toward GDP of countries like India, Bangladesh, and China and also provides livelihood and employment to the field workforce. In India, 58% of the population depends on agriculture, while in Bangladesh, the sector contributes over 11% to GDP and employs more than 37% of the workforce.

Despite this essentiality, plant diseases and post-harvest defects account for over one-third of global agricultural losses, severely affecting various high-value crops.

Fruits occupy a particularly sensitive position in this value chain. Because they are perishable, the market value of a fruit is effectively fixed at the point of inspection by three factors: its species, its current health status, and the extent of visible and morphological surface damage such as bruising, lesions, or rot. A defective unit that goes undetected can accelerate spoilage of neighbouring produce during storage and transport, whereas a sound unit that is wrongly rejected is an avoidable loss to the grower.

Accurate, consistent, and rapid grading is therefore central to reducing post-harvest losses, achieving fair pricing, and redirecting lower-grade produce towards secondary uses such as processing, animal feed, or composting rather than waste.

Traditional disease detection and quality grading rely on manual inspection—slow, subjective, and impractical at scale. Earlier technological alternatives such as hyperspectral and multispectral imaging improved detection capability but were prohibitively costly and lacked real-time performance. Conventional machine vision approaches using handcrafted features and classifiers such as SVM and K-means similarly struggled with generalization under environmental variability.

Recent advances in deep learning—particularly CNN-based architectures including ResNet, EfficientNet, and YOLO augmented with attention mechanisms—have substantially improved automated crop inspection through robust feature extraction, real-time detection, and accurate disease localization. However, much of this work concentrates only on leaf-level disease recognition for a small number of crops and treats classification, localization, and quality grading as separate problems. Studies that do address harvested produce tend to focus on a single species or a single quality attribute, and, to the best of our knowledge, none converts the inspection outcome into a nutritional and economic valuation that supply-chain actors can act upon.

To address this gap, this paper proposes MACISF, a multi-stage software pipeline that integrates crop classification, disease detection, quality grading, and nutritional and economic valuation in a single end-to-end workflow. As illustrated in Figure 1, a fruit image is progressively mapped to its species, its health status, its defect severity grade, and finally a nutritional profile and price estimate. The framework is developed and evaluated on seven economically significant fruit species. The hardware integration of this pipeline, including edge deployment and conveyor-based sorting, is reported separately.

**Figure 1:**
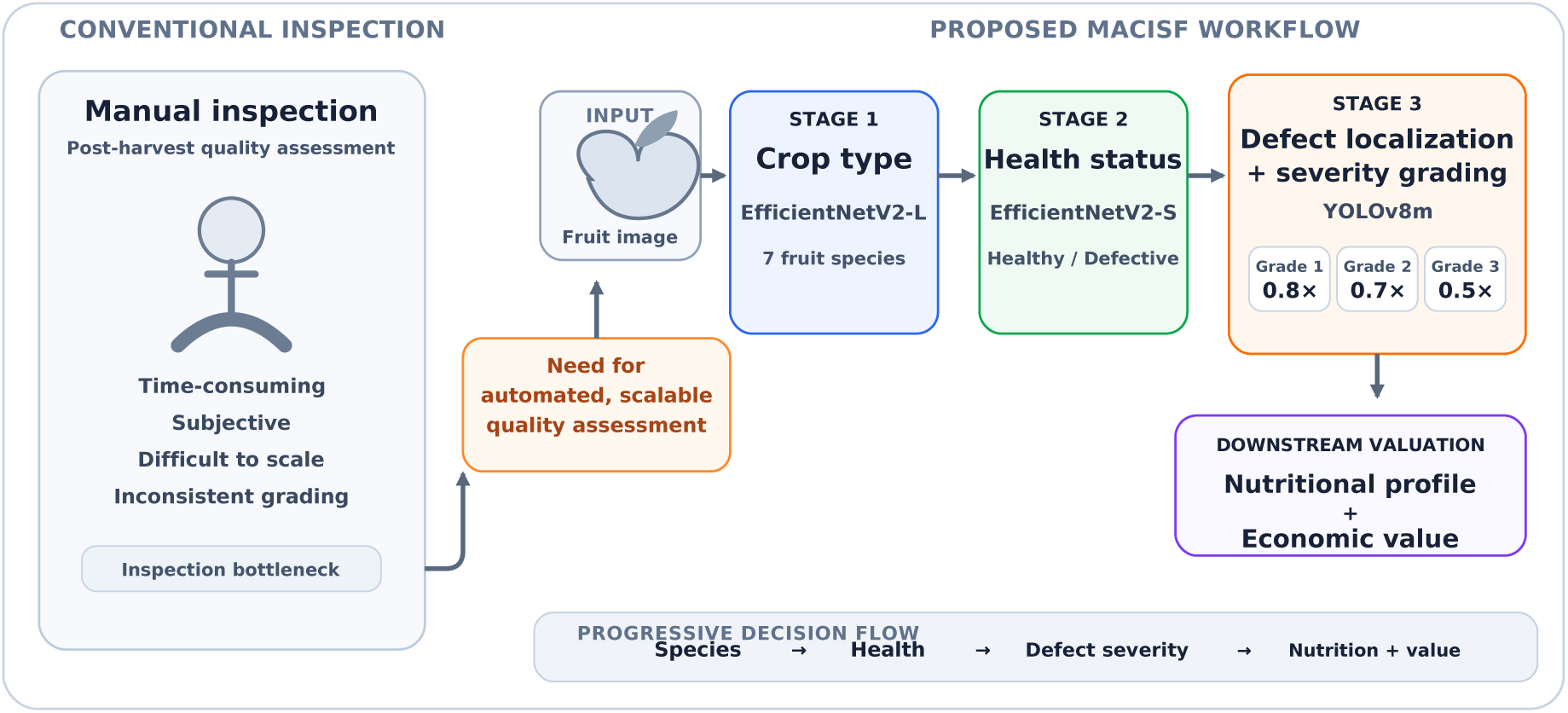
Overview of the post-harvest fruit inspection problem and the proposed MACISF workflow. An input fruit image is progressively mapped to crop type, health status, defect grade, nutritional profile, and economic value.

The main contributions of this work are summarized as follows:

- A unified three-stage software pipeline that integrates crop type classification, species-specific health state detection, defect localization with severity grading, and rule-based nutritional and economic valuation.
- A three-stage progressive fine-tuning protocol for EfficientNetV2-L, combining differential learning rates with Squeeze-Excitation (SE) block calibration, which reaches 99.23% crop type accuracy across seven fruit species.
- Seven species-specific EfficientNetV2-S binary health classifiers trained with a class-weighted loss, reaching 99.39% mean accuracy and 99.51% mean recall on the defective class.
- A three-tier defect severity scheme with a custom bounding-box annotation set and a YOLOv8m grading module whose output is mapped to price multipliers of 0.8×, 0.7×, and 0.5× the healthy baseline.
- A comparative evaluation against CNN and YOLO-family baselines, together with an ablation study of the fine-tuning strategy, label smoothing, augmentation, and SE attention.

The remainder of the paper is organized as follows. Section 2 reviews related work. Section 3 identifies the research gaps and states the problem addressed. Section 4 describes the proposed methodology and the dataset. Section 5 presents the experimental results, comparisons, and ablation study. Section 6 concludes the paper and outlines future scope.

## 2 Related Work

This section reviews prior work in four areas that MACISF draws upon: deep learning for crop disease detection, attention mechanisms and multi-scale feature fusion, object detection and segmentation frameworks, and crop quality grading. Table 1, Table 2, and Table 3 summarize the representative studies in each area.

**Table 1:** Summary of representative deep learning studies on crop disease detection and classification.

| Study | Crop | Method | Key contribution / reported outcome |
| --- | --- | --- | --- |
| Agarwal et al. [1] | Tomato | ToLeD, custom CNN | Trained on PlantVillage (54,305 images, 38 classes); 91.2% accuracy |
| Alhammad et al. [2] | Potato | DL-XAI framework | Leaf disease classification with Grad-CAM explainability |
| Catal Reis and Turk [4] | Potato | MSSCIRNet: depthwise separable convolution and multi-head attention | Competitive performance with reduced parameter count |
| Trivedi et al. [15] | Tomato | Contrast enhancement, grayscale conversion, region-based segmentation | Detection under real (non-laboratory) imaging environments |
| Wang and Liu [16] | Tomato | TomatoDet: YOLOv8 with Swin-DDETR attention | Greenhouse detection under complex lighting |
| Kokate et al. [10] | Tomato | Resource-efficient custom CNN | Suitable for embedded agricultural deployment |

**Table 2:** Summary of representative studies on attention mechanisms, object detection, and segmentation for crop inspection.

| Study | Crop | Method | Key contribution / reported outcome |
| --- | --- | --- | --- |
| Wang et al. [17] | Tomato | CBAM in a YOLOv6 backbone with BiRepGFPN multi-scale fusion | 93.8% mAP on the PlantDoc dataset |
| Khan et al. [8] | Tomato | TomFormer (transformer) with the Hello Stretch robot | Automated image acquisition; long-range dependency modelling of disease symptoms |
| Uddin et al. [28] | Cauliflower | CauliDet: modified YOLOv8 | Localization of three diseases on the VegNet dataset |
| Hu et al. [7] | Apple | ASDINet: few-shot semantic segmentation with global decision and domain attention modules | Apple surface defect localization; small lesions identified as the main bottleneck |
| Chen et al. [5] | General | Survey of detection architectures (sliding window, R-CNN, YOLO, SSD, transformers) | Review of 2D and 3D object detection from images |
| You et al. [18] | Multiple | SimAM-EfficientNet; GP-MI-FGSM adversarial attack | 99.6% accuracy on a multi-class plant disease dataset; robustness evaluation |

**Table 3:** Summary of representative studies on crop quality grading and assessment.

| Study | Crop | Method | Key contribution / reported outcome |
| --- | --- | --- | --- |
| Sharma and Grover [14] | Potato | Hybrid CNN-SVM model | 96.4% precision across four quality classes |
| Kurade et al. [11] | Rice | Image processing module on a Raspberry Pi platform | Low-cost (\$50) automated grain quality evaluation |
| McDonald et al. [12] | Field pea | LDA on 2D color and shape features plus laser-measured 3D seed height | Significantly improved grading accuracy for market grades |

### 2.1 Deep Learning for Crop Disease Detection

Agarwal et al. [1] proposed ToLeD, a custom CNN trained on PlantVillage’s 54,305 RGB images across 38 disease classes, achieving 91.2% accuracy. Alhammad et al. [2] developed a DL-XAI framework for potato leaf disease classification with Grad-CAM explainability. Catal Reis and Turk [4] proposed MSSCIRNet using depthwise separable convolutions and multi-head attention, achieving competitive performance with reduced parameter counts. Trivedi et al. [15] addressed real-environment tomato detection via contrast enhancement, grayscale conversion, and region-based segmentation. Wang and Liu [16] proposed TomatoDet, fusing YOLOv8 with Swin-DDETR attention for greenhouse detection under complex lighting. Kokate et al. [10] contributed a resource-efficient CNN for embedded agricultural deployment.

### 2.2 Attention Mechanisms and Multi-Scale Feature Fusion

Wang et al. [17] integrated the Convolutional Block Attention Module (CBAM) into a YOLOv6 backbone with BiRepGFPN-based multi-scale fusion, achieving 93.8% mAP on the PlantDoc dataset. Khan et al. [8] proposed TomFormer, a transformer architecture paired with the Hello Stretch robotic system for automated image acquisition and long-range dependency modelling of tomato disease symptoms.

### 2.3 Object Detection and Segmentation Frameworks

Uddin et al. [28] developed CauliDet, a modified YOLOv8 model for localization of three cauliflower diseases from the VegNet dataset. Hu et al. [7] proposed ASDINet, a few-shot semantic segmentation network for apple surface defect localization incorporating global decision and domain attention modules, identifying small lesion detection as its primary performance bottleneck. Chen et al. [5] surveyed detection architectures from sliding window methods through R-CNN, YOLO, and SSD series to transformer-based frameworks. You et al. [18] proposed SimAM-EfficientNet achieving 99.6% accuracy on a multi-class plant disease dataset and introduced GP-MI-FGSM, a novel adversarial attack algorithm for robustness evaluation.

### 2.4 Crop Quality Grading and Assessment

Sharma and Grover [14] proposed a hybrid CNN-SVM model for potato quality classification achieving 96.4% precision across four quality classes. Kurade et al. [11] demonstrated low-cost automated rice grain quality evaluation on a Raspberry Pi platform costing $50. McDonald et al. [12] showed that incorporating laser-measured 3D seed height data alongside 2D color and shape features significantly improved LDA grading accuracy for field pea market grades.

## 3 Research Gap and Problem Statement

Despite significant progress across individual components, no existing work, to the best of our knowledge, unifies disease detection, quality grading, and nutritional estimation into a single end-to-end pipeline. Existing studies remain single-task in scope, addressing single crop types or single quality attributes [7, 14, 2]. The critical integration of disease classification, nutritional quality estimation, and market-grade assessment enabling complete economic valuation at the point of inspection remains unaddressed.

### 3.1 Limitations of Existing Studies

Table 4 compares the capabilities reported by the studies reviewed in Section 2 with those of MACISF. Most studies provide a single capability: disease identification, defect localization, or quality grading. None reports nutritional estimation or economic valuation.

**Table 4:** Capability comparison between the reviewed studies and MACISF. A check mark indicates that the capability is reported in the cited work; a dash indicates that it is not reported.

| Study | Disease /<br>defect iden-<br>tification | Defect<br>localization | Quality<br>grading | Nutritional<br>estimation | Economic<br>valuation |
| --- | --- | --- | --- | --- | --- |
| Agarwal et al. [1] | ✓ | – | – | – | – |
| Alhammad et al. [2] | ✓ | – | – | – | – |
| Catal Reis and Turk [4] | ✓ | – | – | – | – |
| Trivedi et al. [15] | ✓ | – | – | – | – |
| Kokate et al. [10] | ✓ | – | – | – | – |
| Khan et al. [8] | ✓ | – | – | – | – |
| You et al. [18] | ✓ | – | – | – | – |
| Wang and Liu [16] | ✓ | ✓ | – | – | – |
| Wang et al. [17] | ✓ | ✓ | – | – | – |
| Uddin et al. [28] | ✓ | ✓ | – | – | – |
| Hu et al. [7] | ✓ | ✓ | – | – | – |
| Sharma and Grover [14] | – | – | ✓ | – | – |
| Kurade et al. [11] | – | – | ✓ | – | – |
| McDonald et al. [12] | ✓ | – | ✓ | – | – |
| **MACISF (proposed)** | ✓ | ✓ | ✓ | ✓ | ✓ |

The following research gaps are identified:

1. **Fragmented tasks.** Disease identification, defect localization, and quality grading are addressed in isolation, so no single system produces a complete quality profile of an inspected unit.
2. **Narrow crop scope.** Leaf-level studies concentrate on tomato and potato, and grading studies typically address a single commodity [14, 12].
3. **No severity-aware grading with localization.** Defect localization frameworks do not assign market grades [7], while grading models do not localize the defects that determine the grade [14].
4. **No annotated defect-severity data.** No publicly available dataset provides bounding-box annotations of fruit surface defects classified by severity.
5. **No downstream valuation.** Inspection outcomes are not linked to nutritional impact or to economic value at the point of inspection.
6. **Robustness to imaging variability.** Laboratory datasets with uniform backgrounds transfer poorly to real conditions, and small, low-contrast lesions remain the main bottleneck for detection [7, 28].

### 3.2 Problem Statement and Objectives

Given an RGB image of a single fruit, the problem addressed in this paper is to determine automatically its crop species, its health status, and, if defective, the location and severity of its defects, and to translate this outcome into a nutritional and economic valuation. The specific objectives are:

1. to classify the crop type among seven fruit species;
2. to determine the healthy or defective state of the specimen using species-specific classifiers;
3. to localize defects and assign a three-tier severity grade;
4. to map species, health status, and grade to a nutritional profile and a per-unit price estimate; and
5. to evaluate the pipeline against established baselines and quantify the contribution of each design component through an ablation study.

### 3.3 How MACISF Addresses the Gaps

Table 5 maps each identified gap to the corresponding design decision in MACISF.

**Table 5:** Research gaps and the corresponding MACISF design responses.

| Research gap | MACISF design response |
| --- | --- |
| Fragmented tasks | Single sequential pipeline: crop type → health state → defect grade → valuation |
| Narrow crop scope | Seven fruit species with modular, species-specific Stage 2 classifiers |
| No severity-aware grading with localization | YOLOv8m jointly localizes defects and assigns one of three severity grades |
| No annotated defect-severity data | Custom bounding-box annotation of defects with three grade labels (LabelImg) |
| No downstream valuation | Rule-based nutritional profile and price multipliers of $0.8\times$, $0.7\times$, and $0.5\times$ |
| Robustness to imaging variability | Extensive augmentation, Mixup/CutMix, and test-time augmentation |

## 4 Proposed Methodology and Dataset

### 4.1 Overview of the Proposed Framework

The proposed Multistage AI Crop Inspection and Segmentation Framework (MACISF) is designed as a sequential, multi-stage intelligent pipeline that processes fruit images through a series of specialized deep learning modules, each building upon the output of the preceding stage to progressively generate a complete quality profile—from crop identification to disease status to market grade and final nutritional and economic valuation. As illustrated in Figure 3, the pipeline comprises a two-stage classification system built upon EfficientNetV2 and a grade-level object detection module built upon YOLOv8 with custom annotations. The overall design architecture of MACISF prioritizes practical deployability alongside classification accuracy, recognizing that a system destined for real agricultural supply chain integration must operate effectively under variable lighting conditions and surface occlusion.

**Figure 2:**
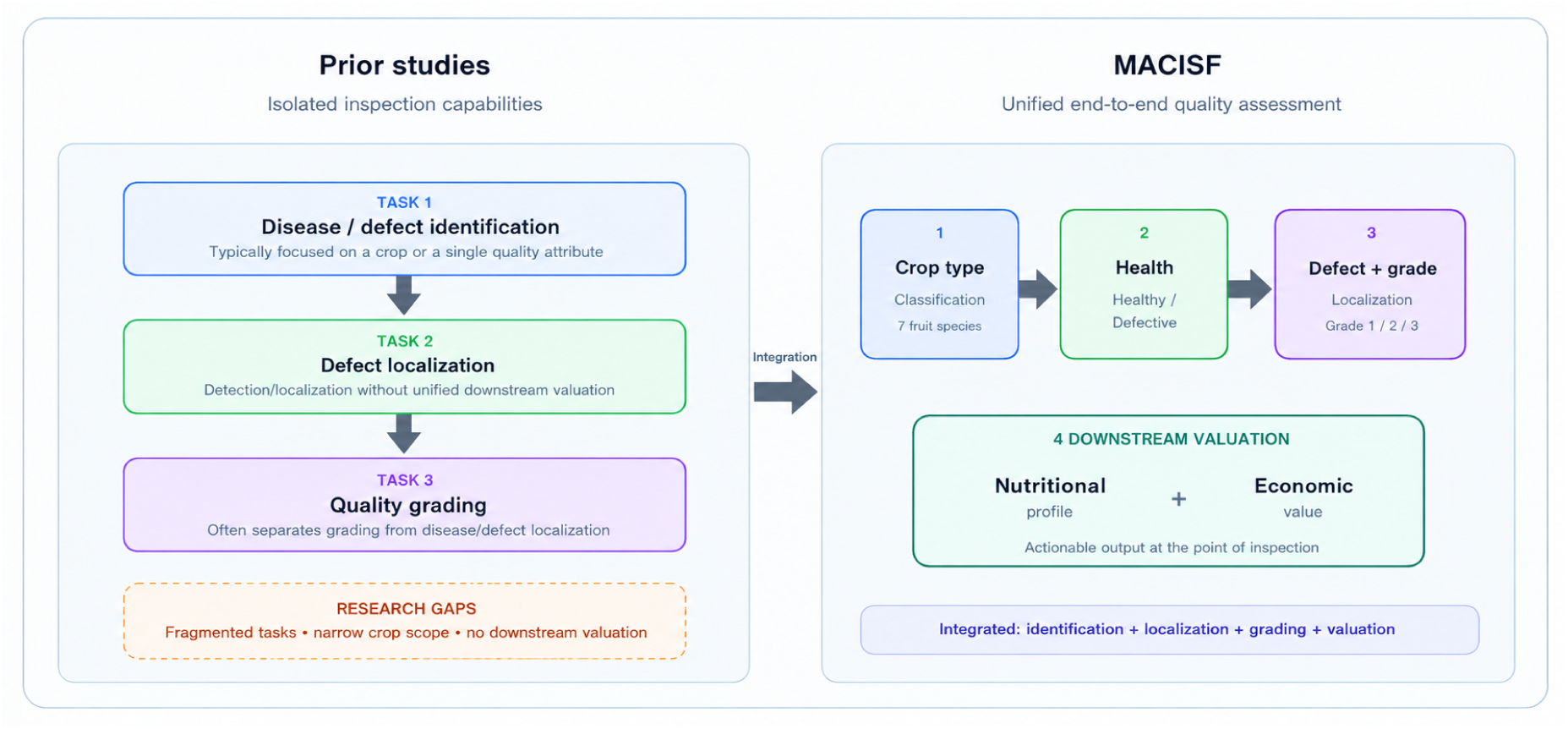
Positioning of MACISF relative to existing work: prior studies address isolated tasks, whereas MACISF integrates classification, localization, grading, and valuation in one pipeline.

**Figure 3:**
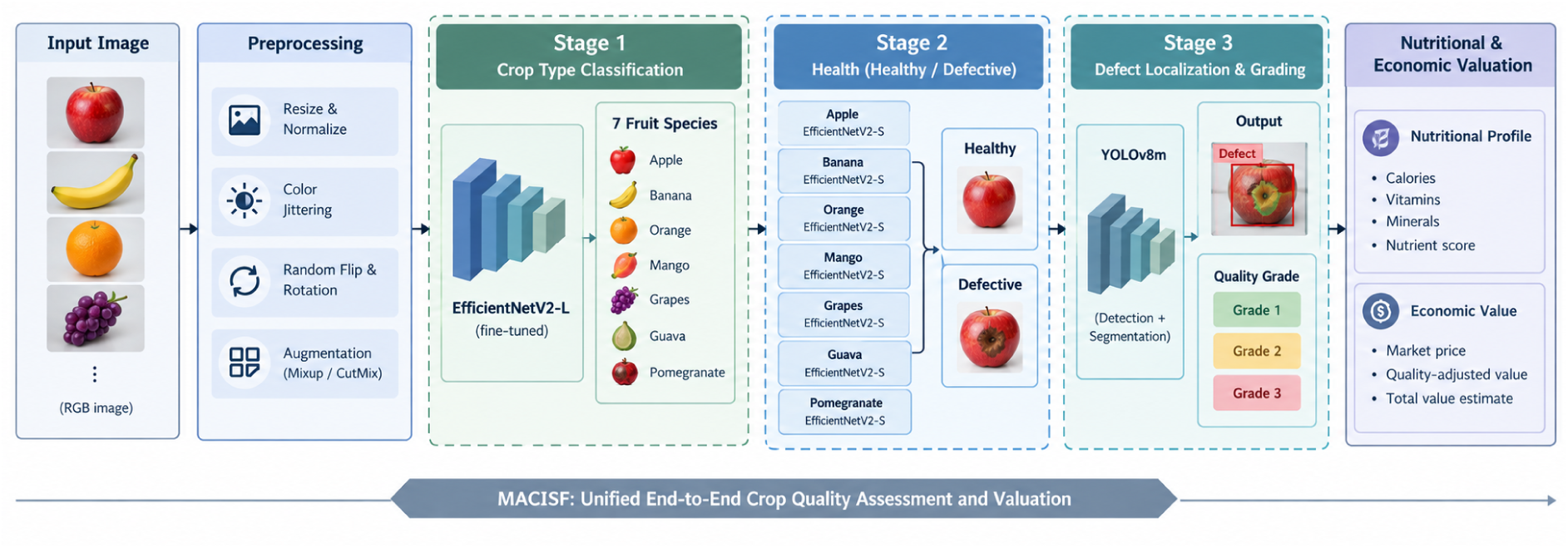
Model architecture of the proposed MACISF pipeline. The framework processes an input fruit image through preprocessing, Stage 1 EfficientNetV2-L crop type classification, Stage 2 species-specific EfficientNetV2-S health detection, Stage 3 YOLOv8m defect localization and quality grading, and finally nutritional and economic valuation.

The framework currently covers seven economically significant fruit crops: apple (*Malus domestica*), lime (*Citrus aurantiifolia*), banana (*Musa* spp.), mango (*Mangifera indica*), guava (*Psidium guajava*), pomegranate (*Punica granatum*), and orange (*Citrus sinensis*). These seven crops were selected on the basis of their combined market significance in South Asian agricultural economies, their high susceptibility to post-harvest surface defects and disease-related quality degradation, and the availability of sufficiently diverse image data across healthy and defective classes to support robust deep learning training.

### 4.2 Stage 1 — Crop Type Classification Using EfficientNetV2-L

Stage 1 of the MACISF pipeline identifies the crop species from the seven target classes using a single EfficientNetV2-L backbone with one classification head. The predicted species determines which of the seven species-specific Stage 2 health classifiers receives the image. Identifying the species first, rather than training one shared health classifier for all species, allows each Stage 2 model to specialise in the defect presentations of a single fruit.

#### 4.2.1 Architecture Selection

EfficientNetV2-L was selected as the backbone architecture on the basis of its demonstrated efficiency in classification accuracy relative to prior-generation EfficientNetV1 and contemporaneous architectures such as ResNet-101 and DenseNet-201. Proposed by Tan and Le [19], EfficientNetV2 improves upon V1 through three principal innovations: substitution of Fused-MBConv layers in early network stages—replacing depthwise separable convolutions with full convolutions fused with the projection layer for faster GPU training; a progressive training strategy that dynamically adjusts image resolution and regularization strength during pre-training; and a Neural Architecture Search procedure optimized jointly for training speed and parameter efficiency. The V2-L variant achieves an ImageNet-1k Top-1 accuracy of 85.8%, compared to 77.4% for ResNet-101 and 77.3% for DenseNet-201—margins of 8.4 and 8.5 percentage points respectively over architectures that would otherwise represent strong baselines for fine-grained visual classification (Table 6).

**Table 6:** Comparison of candidate backbone architectures. Parameter counts are the published values for each architecture with its standard 1000-class ImageNet head.

| Architecture | ImageNet<br>Top-1 (%) | Feature Dim | Parameters<br>(M) | SE Attention |
| --- | --- | --- | --- | --- |
| EfficientNet-V2-L (selected) | 85.8 | 1280 | 118.5 | Yes (stages 4–6) |
| ResNet-101 | 77.4 | 2048 | 44.5 | No |
| DenseNet-201 | 77.3 | 1920 | 20.0 | No |

The EfficientNetV2-L feature extractor is organized into eight progressive stages. Stages 1–3 (`features[1]–features[3]`) employ Fused-MBConv blocks with 32, 64, and 96 channels respectively, extracting foundational low-level signals including edges, color gradients, surface textures, and basic structural shapes. Stages 4–6 (`features[4]–features[6]`) transition to standard MBConv blocks augmented with Squeeze-Excitation (SE) channel attention, expanding representations from 96 to 384 dimensions. The SE mechanism compresses each feature channel to a global summary activation and learns a reweighting scheme that automatically assigns greater representational importance to diagnostically relevant visual signals—tonal gradients of ripe mango, waxy surface reflectance of healthy lime, or darkened tissue patterns of bruised apple—without requiring explicit human guidance. Stage 7 (`features[7]`) projects the 384-dimensional output to 1280 dimensions, which Adaptive Average Pooling collapses into the final compact feature vector used by the classification head.

The classification head follows a three-layer architecture: Linear(1280 → 640) → BatchNorm → SiLU → Dropout(0.4) → Linear(640 → 320) → BatchNorm → SiLU → Dropout(0.2) → Linear(320 → 7) output logits. SiLU activation was selected to match the internal activation function of the EfficientNetV2-L backbone, ensuring activation distribution consistency at the backbone–head interface. Raw logits are passed directly to the loss function to maintain numerical stability under mixed-precision training.

#### 4.2.2 Transfer Learning and Fine-Tuning Strategy

The EfficientNetV2-L backbone was initialized with ImageNet-1k pretrained weights. A three-stage progressive fine-tuning strategy was adopted to balance knowledge transfer from ImageNet pre-training against domain adaptation to the agricultural fruit classification task. The additional fine-tuning stage relative to standard two-phase approaches is necessitated by the greater depth of EfficientNetV2-L compared to ResNet or DenseNet equivalents, requiring more careful incremental unfreezing to avoid destabilizing SE attention patterns acquired during pre-training.

##### Stage 1 — Head Warm-Up (15 epochs)

All backbone parameters frozen; only the classification head trained. AdamW optimizer (LR = 3 × 10*^−^*^4^, WD = 1 × 10*^−^*^4^) with CosineAn-nealingWarmRestarts (*T*_0_ = 15, min LR = 1 × 10*^−^*^6^). This phase allows randomly initialized head weights to converge to a stable parameter regime before backbone parameters are updated, preventing large head-layer gradients from distorting pre-trained feature representations on the first forward pass.

##### Stage 2 — Deep Feature Adaptation (20 epochs)

Backbone stages `features[5]`, `features[6]`, and `features[7]` unfrozen; `features[0]–features[4]` remain frozen. AdamW reconfigured with differential learning rates: 3 × 10*^−^*^4^ for the head and 5 × 10*^−^*^6^ for unfrozen backbone blocks. CosineAnnealingWarmRestarts (*T*_0_ = 5, *T*_mult_ = 2, min LR = 1 × 10*^−^*^7^). The unfrozen stages encompass the deepest MBConv+SE attention blocks and head convolution—those layers whose activations are most task-specific and most likely to benefit from agricultural domain adaptation.

##### Stage 3 — SE Block Calibration (15 epochs)

`features[4]` additionally unfrozen at a conservative LR of 5 × 10*^−^*^7^—one order of magnitude below Stage 2’s backbone rate. This asymmetry is intentional: `features[4]` contains SE attention blocks encoding general-purpose channel weighting relationships from ImageNet, and the low learning rate provides a gentle recalibration toward fruit-domain color-channel importance distributions without erasing the underlying attention structure. CosineAnnealingWarmRestarts (*T*_0_ = 5, *T*_mult_ = 2, min LR = 1 × 10*^−^*^8^).

Across all three stages: AdamW (WD = 1 × 10*^−^*^4^), gradient clipping (max norm = 1.0), label smoothing (*ε* = 0.1) on the cross-entropy loss, AMP training with GradScaler, and early stopping (patience = 6 epochs, min delta = 1 × 10*^−^*^4^) on validation accuracy with best-checkpoint saving.

#### 4.2.3 Input Preprocessing

During evaluation and inference, all input images were resized to 384 × 384 pixels using bilinear interpolation, matching the input resolution used during training. Pixel values were normalized to the range [0, 1] and standardized using the ImageNet channel-wise mean [0.485, 0.456, 0.406] and standard deviation [0.229, 0.224, 0.225].

During training, an extensive online data augmentation pipeline was applied to each batch prior to normalisation to improve model robustness to the imaging variability characteristic of field and conveyor belt deployment conditions. The following augmentation operations were applied sequentially:

- **RandomResizedCrop** — output size 384 × 384; scale range (0.50, 1.00); aspect ratio range (0.75, 1.33)
- **RandomHorizontalFlip** — probability 0.50
- **RandomVerticalFlip** — probability 0.20
- **RandomRotation** — range ±35*^◦^*
- **ColorJitter** — brightness ±0.6; contrast ±0.6; saturation ±0.5; hue ±0.15; applied with probability 0.85
- **RandomGrayscale** — probability 0.05
- **RandomPerspective** — distortion scale 0.25; probability 0.30
- **GaussianBlur** — kernel 5 × 5; sigma uniformly sampled from (0.1, 2.5); probability 0.30
- **RandomSolarize** — threshold 110; probability 0.12
- **RandomAutocontrast** — probability 0.20
- **RandomEqualize** — probability 0.12
- **RandomErasing** — probability 0.45; patch area scale (0.02, 0.28); aspect ratio range (0.30, 3.30); fill values randomised

RandomErasing was specifically included to simulate partial occlusion of fruit surfaces by leaves, packaging fragments, or adjacent produce on the conveyor belt.

In addition to the per-image augmentation pipeline, batch-level augmentation was applied during training using a stochastic combination of Mixup (*α* = 0.4) and CutMix (*α* = 1.0), with each mini-batch randomly assigned to one of the two strategies with equal probability. Both strategies produce convex combinations of training examples and their labels, regularising the model against overconfident boundary decisions. For CutMix, rectangular patches of area proportional to (1 − *λ*), where *λ* ∼ Beta(1.0, 1.0), were replaced with the corresponding patch from a randomly selected batch partner, with labels mixed in proportion to the actual pixel area ratio.

#### 4.2.4 Test-Time Augmentation

At inference time, Test-Time Augmentation was applied to improve the prediction robustness of the model by averaging the softmax probability outputs across eight deterministic augmentation variants of each input image. The eight variants comprise: the unmodified image, horizontal flip, vertical flip, 90*^◦^* rotation, 180*^◦^* rotation, 270*^◦^* rotation, combined horizontal flip with 90*^◦^* rotation, and brightness increase by a factor of 1.2. Softmax probability vectors from all eight variants were averaged, and the final prediction was obtained by taking the argmax of the averaged distribution.

#### 4.2.5 Loss Function and Model Selection

The Stage 1 training loss is the cross-entropy over the seven crop classes with label smoothing of 0.1:

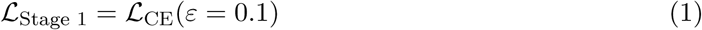

Validation accuracy on the held-out validation set was used as the model selection criterion for early stopping and checkpoint saving throughout all three fine-tuning stages.

### 4.3 Stage 2 — Health State Classification Using Species-Specific EfficientNetV2-S Classifiers

Once the crop type has been determined, the image is passed to the health state classifier of that species, which decides whether the specimen is healthy or defective. Only specimens classified as defective proceed to Stage 3.

#### 4.3.1 Species-Conditioned Classification Architecture

Stage 2 consists of seven independent binary classifiers, one per fruit species. Each classifier uses an EfficientNetV2-S backbone initialized with ImageNet-1k pretrained weights and is trained exclusively on the healthy and defective images of its own species. The classification head has the same layer structure as the Stage 1 head, ending in a two-way output layer, Linear(320 → 2). Input resolution, preprocessing, augmentation, and test-time augmentation follow the Stage 1 configuration (Section 4.2.3 and Section 4.2.4). Each classifier is fine-tuned in two stages (Table 9): a 10-epoch head warm-up with the backbone frozen, followed by 20 epochs of full fine-tuning with the backbone at a learning rate of 1 × 10*^−^*^5^ and the head at 3 × 10*^−^*^4^. Because each classifier sees only one species, the defect presentations specific to that species are modelled without interference from the other fruit classes, and a new species can be added by training one additional classifier.

#### 4.3.2 Class-Weighted Loss for Class Imbalance

Healthy and defective images are nearly balanced in every species (Table 7); the most skewed species, guava, has 1716 healthy and 2021 defective images. To guard against the residual skew, the cross-entropy loss of each Stage 2 classifier is weighted per class, with weights inversely proportional to the class frequency in the training split, in addition to label smoothing of 0.1.

**Table 7:** Class distribution across the seven fruit species in the full fruit dataset, before the 70:15:15 training/validation/test split.

| Fruit Species | Healthy Images | Defective Images | Total |
| --- | --- | --- | --- |
| Apple | 1903 | 1878 | 3781 |
| Mango | 2100 | 1976 | 4076 |
| Banana | 1886 | 1932 | 3818 |
| Orange | 1615 | 1538 | 3153 |
| Guava | 1716 | 2021 | 3737 |
| Lime | 2063 | 2100 | 4163 |
| Pomegranate | 2083 | 1951 | 4034 |
| **Total** | **13366** | **13396** | **26762** |

### 4.4 Stage 3 — Quality Grading Using YOLOv8 with Custom Annotations

Following the determination of crop species and health status, produce classified as defective is passed to the third stage of the pipeline: a quality grading and defect localization module based on the YOLOv8 object detection architecture with a fully custom annotation scheme developed specifically for this study.

#### 4.4.1 Grading Scheme Definition

A three-tier grading scheme classifies defective produce by the nature, extent, and distribution of surface damage, each tier carrying distinct implications for market value, nutritional profile, and appropriate downstream use:

- **Grade 1 (Minor Defect):** Small, localized surface blemishes, minor bruising, or early-stage lesions covering less than 10% of the total visible surface area. Nutritional value is not meaningfully compromised, and affected regions can typically be removed during preparation with minimal waste. Suitable for direct retail at a marginal price reduction. Price multiplier: 0.8× the species-specific healthy baseline market rate.
- **Grade 2 (Moderate Defect):** Visibly enlarged lesions, moderate bruising, or multi-focal surface damage covering 10–30% of the visible surface area. Retains significant nutritional content but is unsuitable for premium retail; suitable for secondary processing, juice extraction, or controlled-atmosphere storage. Price multiplier: 0.7× the healthy baseline rate.
- **Grade 3 (Severe Defect):** Extensive surface damage, deep lesions, advanced rot, or disease spread covering more than 30% of the visible surface area. Significant nutritional degradation and contamination risk render the produce unsuitable for direct retail; redirected to composting, animal feed, or biogas production pipelines. Price multiplier: 0.5× the healthy baseline rate.

These scalar multipliers enable MACISF to generate an immediate per-unit economic valuation at the point of inspection, transforming classification output into an actionable financial signal communicated directly to supply chain management systems and farmer-facing dashboards (Figure 4).

**Figure 4:**
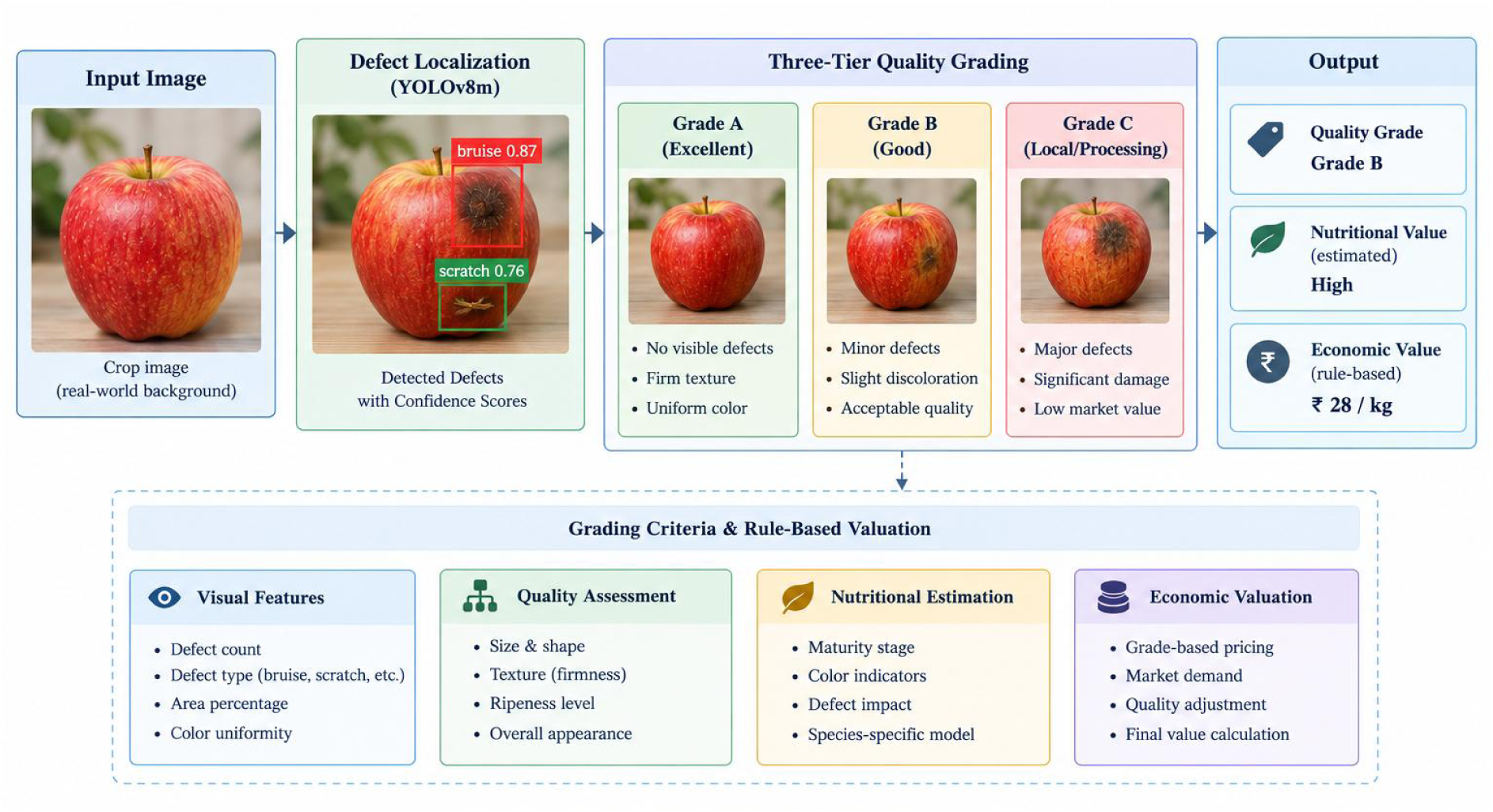
Illustration of the three-tier defect grading scheme with the surface-damage thresholds and price multipliers (0.8×, 0.7×, 0.5×) assigned to Grades 1, 2, and 3.

#### 4.4.2 YOLOv8 Architecture and Configuration

YOLOv8, a widely adopted member of the You Only Look Once real-time object detection family, was selected as the grading module architecture on the basis of its strong balance between detection speed and bounding box localization accuracy and its native support for instance segmentation alongside bounding box detection. The YOLOv8m (medium) variant was selected, providing a strong balance between the parameter count and inference performance.

The grading module was configured to perform simultaneous defect localization and grade classification: for each input image of a defective produce specimen, the YOLOv8 model outputs one or more bounding boxes enclosing detected defect regions, each annotated with a class label corresponding to one of the three grade-defining defect severity categories. The final grade assigned to the specimen is determined by the maximum severity class detected across all the predicted bounding boxes.

#### 4.4.3 Custom Annotation Process

Given that no existing publicly available dataset provides bounding box-level annotations of fruit surface defects classified according to the three-tier grading scheme defined for this study, a dedicated custom annotation effort was undertaken specifically for this work. Fruit images from the composite dataset were annotated using the LabelImg annotation tool, with annotators drawing tight bounding boxes around each visible defect region on the fruit surface and assigning one of three grade labels based on the defect severity criteria defined in Section 4.4.1.

### 4.5 Nutritional Value Estimation and Economic Valuation Module

Downstream of the grading classification, MACISF incorporates a rule-based nutritional value estimation and economic valuation module that maps the detected species, health status, and grade label to a structured output profile which is communicated to any connected digital reporting interface.

For each inspected fruit unit, the valuation module retrieves the species-specific baseline nutritional profile—expressed in terms of key parameters including the caloric content, Vitamin C concentration, dietary fibre content, sugar concentration, and water activity. For Grade 1 produce, the full baseline nutritional profile is retained with no modifications, as minor surface defects do not significantly alter the nutritional composition of the unaffected flesh. For Grade 2 produce, a moderate reduction coefficient is applied to vitamin content and antioxidant values, reflecting the documented tendency of moderate fungal and bacterial infections to accelerate oxidative degradation of heat-sensitive micronutrients in adjacent tissue. For Grade 3 produce, substantial reduction coefficients are applied across most nutritional parameters, reflecting the tissue compromise associated with severe surface damage and advanced pathogen colonization.

### 4.6 Dataset Composition

The composite dataset assembled for training and evaluation of the MACISF pipeline integrates multiple publicly available benchmark datasets to achieve the breadth of species coverage, defect diversity, and environmental variability required for a practically deployable multi-crop inspection system.

#### 4.6.1 Public Benchmark Datasets

The following publicly available datasets form the foundation of the training corpus:

- **PlantVillage Dataset** [20]: 54,305 RGB images across 38 disease and healthy classes covering tomato, potato, bell pepper, and other species, captured under controlled laboratory conditions. Despite limited real-world transferability due to uniform backgrounds, it provides high-quality, consistently labeled disease symptom presentations—particularly valuable for tomato and potato subsets contributing to Stage 2 health classification training.
- **Plant Disease Dataset**: 79,265 images across 42 classes covering healthy and diseased tomato, potato, pepper, and other species—the largest labeled disease corpus in the composite training set, offering substantially greater class diversity than PlantVillage for robust Stage 2 classifier training across a wider range of disease presentations.
- **Plant-Doc Dataset** [21]: 2,598 images across 17 disease classes captured in natural outdoor environments with complex, realistic backgrounds. Training on Plant-Doc images substantially improves Stage 2 generalization to uncontrolled imaging conditions.
- **IPM Dataset**: 3,739 images covering various diseases across tomato, potato, and pepper, contributing additional diversity in disease type coverage and imaging conditions.
- **APS Images Dataset**: 7,000 images covering various disease types across tomato, potato, pepper, and other species, providing further diversity in imaging conditions and disease presentations.
- **PLD Dataset** [22]: 4,082 images labeled for early blight, late blight, and healthy potato leaves, providing focused high-quality data for the potato-specific Stage 2 classifier, complementing the broader PlantVillage and Plant Disease Dataset potato subsets.

#### 4.6.2 Fruit Dataset Class Distribution

Table 7 summarizes the class distribution across the seven species in the full fruit dataset (before the 70:15:15 split into training, validation, and test subsets), and Figure 5 shows representative samples.

**Figure 5:**
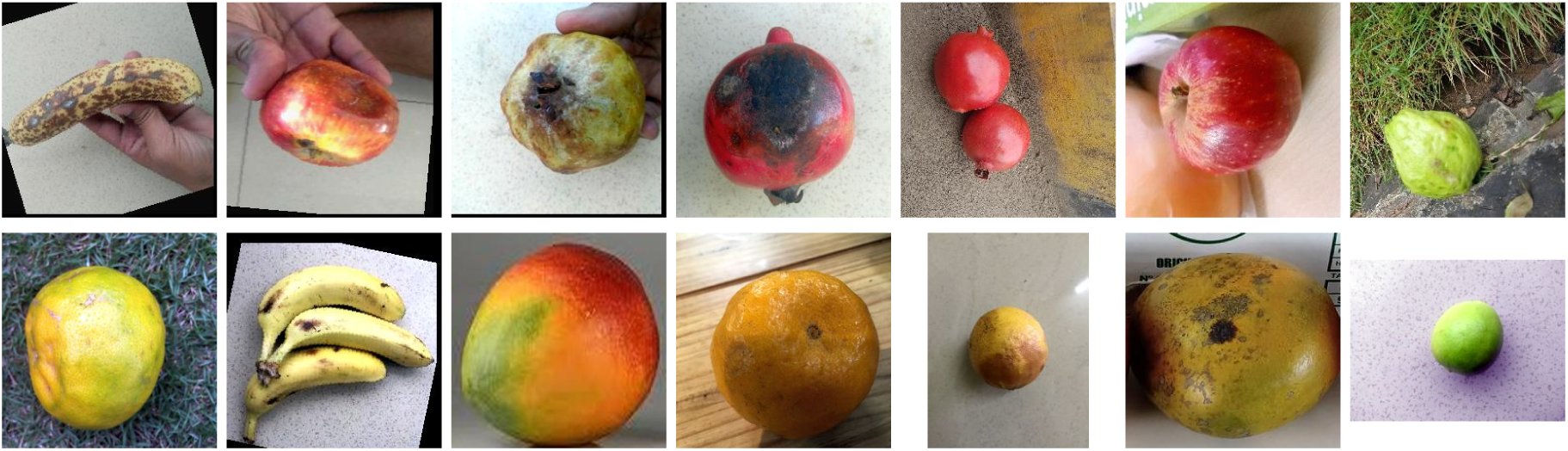
Representative healthy and defective samples from each of the seven fruit species in the composite dataset.

#### 4.6.3 Custom Annotation Dataset for YOLOv8 Grading Module

The grading module training dataset consists of defective fruit images from the entire fruit dataset, subjected to the bounding box annotation process described in Section 4.4.3. Each annotated image contains one or more bounding boxes enclosing visible defect regions, with each box assigned one of the three grade labels (Grade 1, Grade 2, Grade 3) according to the defect severity criteria.

### 4.7 Experimental Setup and Evaluation Metrics

All model training experiments for Stages 1 and 2 were conducted on a workstation equipped with an NVIDIA RTX A5500 GPU with 24 GB VRAM and an Intel Xeon w7-2475X processor, using PyTorch 2.0 with the torchvision EfficientNetV2 implementations. YOLOv8 training for Stage 3 was performed using the Ultralytics YOLOv8 Python package under PyTorch 2.0. Training, validation, and test splits were established at 70:15:15 ratios across all datasets, with stratified splitting applied to preserve class distribution balance across all three subsets.

Model performance across all stages is evaluated using the following metrics: classification accuracy, precision, recall, F1-score, and Area Under the Receiver Operating Characteristic Curve (AUC-ROC) for Stages 1 and 2; and mean Average Precision at IoU threshold 0.5 (mAP@0.5) and mAP@0.5:0.95 for Stage 3 YOLOv8 detection.

## 5 Results and Discussion

This section presents a comprehensive evaluation of the Multistage AI Crop Inspection and Segmentation Framework (MACISF) across all three pipeline stages: crop type classification (Stage 1), health status classification (Stage 2), and quality grade detection (Stage 3). The results are organized as follows: Section 5.1 describes the experimental environment and training configuration; Section 5.2 details the evaluation metrics employed; Section 5.3 presents the Stage 1 crop type classification results; Section 5.4 presents the Stage 2 health status classification results; Section 5.5 evaluates the Stage 3 YOLOv8 grading module; Section 5.6 provides comparative analysis against baselines; and Section 5.7 summarizes the results.

### 5.1 Experimental Environment and Configuration

All the training procedures of the model for Stages 1 and 2 were conducted on a dedicated deep learning workstation, while the YOLOv8 grading module for Stage 3 was trained using the Ultralytics YOLOv8 Python package. Table 8 summarizes the complete hardware and software configuration used across the training and validation phases.

**Table 8:** Hardware and software configuration for training.

| Component | Specification |
| --- | --- |
| GPU (Training) | NVIDIA RTX A5500, 24GB VRAM |
| CPU | Intel(R) Xeon(R) w7-2475X |
| Deep Learning Framework | PyTorch 2.0 |
| EfficientNetV2 Library | torchvision — EfficientNetV2-L and EfficientNetV2-S pretrained on ImageNet-1k |
| YOLOv8 Library | Ultralytics YOLOv8 (Python package, PyTorch backend) |
| Operating System | Ubuntu 20.04 LTS |
| Python Version | Python 3.9 |

Table 9 presents the complete set of hyperparameters and training configuration for the Stage 1 EfficientNetV2-L crop type classifier and the Stage 2 EfficientNetV2-S health state classifiers. The three-stage progressive fine-tuning strategy described in Section 4.2.2 is reflected in the epoch allocation and differential learning rate assignments across stages.

**Table 9:** Hyperparameter configuration for Stage 1 and Stage 2 EfficientNetV2 classifiers.

| Hyperparameter | Stage 1 (Crop Type) | Stage 2 (Health State) |
| --- | --- | --- |
| Architecture | EfficientNetV2-L | EfficientNetV2-S ($\times 7$ species) |
| Input Resolution | $384 \times 384$ px | $384 \times 384$ px |
| Pre-training Dataset | ImageNet-1k | ImageNet-1k |
| Fine-tuning Stages | 3 (Head Warm-Up, Deep Adapt., SE Calib.) | 2 (Head Warm-Up, Full Fine-Tune) |
| Epochs (Stage 1 / 2 / 3) | 15 / 20 / 15 = 50 total | 10 / 20 = 30 total |
| Optimizer | AdamW | AdamW |
| Initial LR (Head) | $3 \times 10^{-4}$ | $3 \times 10^{-4}$ |
| LR (Unfrozen Backbone) | $5 \times 10^{-6}$ / $5 \times 10^{-7}$ | $1 \times 10^{-5}$ |
| Weight Decay | $1 \times 10^{-4}$ | $1 \times 10^{-4}$ |
| LR Scheduler | CosineAnnealingWarmRestarts | CosineAnnealingWarmRestarts |
| Label Smoothing ($\epsilon$) | 0.1 | 0.1 |
| Dropout (Head Layer 1 / 2) | 0.4 / 0.2 | 0.4 / 0.2 |
| Batch Size | 16 | 16 |
| Gradient Clipping | Max norm = 1.0 | Max norm = 1.0 |
| Mixed Precision (AMP) | Enabled (GradScaler) | Enabled (GradScaler) |
| Early Stopping Patience | 6 epochs | 6 epochs |
| Dataset Split | 70 / 15 / 15 (train/val/test) | 70 / 15 / 15 (train/val/test) |

### 5.2 Evaluation Metrics

Model performance across all three pipeline stages is evaluated using a consistent set of quantitative metrics drawn from standard practice in deep learning classification and object detection research. For the classification stages (Stage 1 and Stage 2), the primary evaluation metrics are classification accuracy, precision, recall, F1-score, and the Area Under the Receiver Operating Characteristic Curve (AUC-ROC). For Stage 1, per-class accuracy is computed one-vs-rest, overall accuracy is the fraction of all test images that are classified correctly, and the overall precision, recall, F1-score, and AUC-ROC are macro-averaged, so that all fruit species receive equal weight irrespective of their sample count. For Stage 2, the overall row is the mean over the seven species-specific classifiers. These metrics are defined as follows:

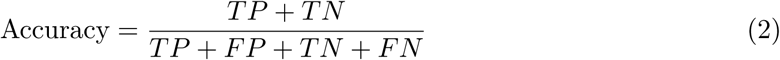

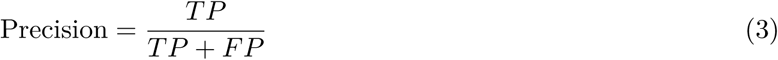

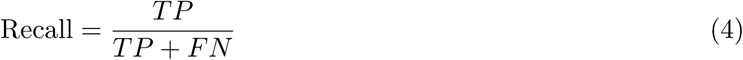

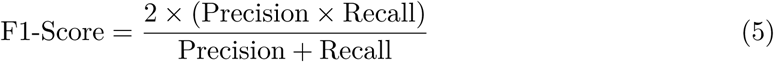

For the Stage 3 YOLOv8 grading module, performance is evaluated using Mean Average Precision at IoU threshold 0.5 (mAP@0.5) and mAP across IoU thresholds from 0.5 to 0.95 in steps of 0.05 (mAP@0.5:0.95). The mAP@0.5 metric measures the model’s ability to correctly detect and classify defect regions with at least 50 percent bounding box overlap with the ground truth annotation, while mAP@0.5:0.95 provides a more stringent assessment of localization precision across a range of overlap thresholds.

### 5.3 Stage 1: Crop Type Classification Results

The Stage 1 EfficientNetV2-L crop type classifier was trained and evaluated on the composite seven-class fruit dataset comprising apple, lime, banana, mango, guava, pomegranate, and orange specimens. The model was trained following the three-stage progressive fine-tuning strategy described in Section 4.2.2, and all results reported in this section correspond to evaluation on the held-out test set (15 percent of the total data) following completion of all three fine-tuning stages (Table 10).

**Table 10:** Per-class and overall performance of the Stage 1 EfficientNetV2-L crop type classifier on the test set. All values are expressed as percentages except AUC-ROC. Per-class accuracy is computed one-vs-rest. In the overall row, accuracy is the fraction of all test images classified correctly, and precision, recall, F1-score, and AUC-ROC are macro averages of the seven species.

| Fruit Species | Accuracy (%) | Precision (%) | Recall (%) | F1-Score (%) | AUC-ROC |
| --- | --- | --- | --- | --- | --- |
| Apple | 99.70 | 98.77 | 99.12 | 98.94 | 0.999 |
| Lime | 99.88 | 99.52 | 99.68 | 99.60 | 0.998 |
| Banana | 100.00 | 100.00 | 100.00 | 100.00 | 1.000 |
| Mango | 99.58 | 99.01 | 98.20 | 98.60 | 0.988 |
| Guava | 99.55 | 98.22 | 98.57 | 98.40 | 0.967 |
| Pomegranate | 99.90 | 99.67 | 99.67 | 99.67 | 0.998 |
| Orange | 99.85 | 99.37 | 99.37 | 99.37 | 1.000 |
| **Macro Avg** | **99.23** | **99.22** | **99.23** | **99.23** | **0.993** |

The Stage 1 model reached an overall classification accuracy of 99.23 percent on the test set, with macro-averaged precision of 99.22 percent, recall of 99.23 percent, F1-score of 99.23 percent, and AUC-ROC of 0.993, demonstrating strong generalization across the seven fruit species. Banana and orange were separated perfectly in the one-vs-rest ROC analysis (AUC-ROC of 1.000), and AUC-ROC exceeded 0.99 for apple, lime, banana, pomegranate, and orange. Mango and guava were the most challenging species: mango had the lowest recall (98.20 percent) and an AUC-ROC of 0.988, while guava had the lowest per-class accuracy (99.55 percent), the lowest precision (98.22 percent), and the lowest AUC-ROC (0.967).

**Figure 6:**
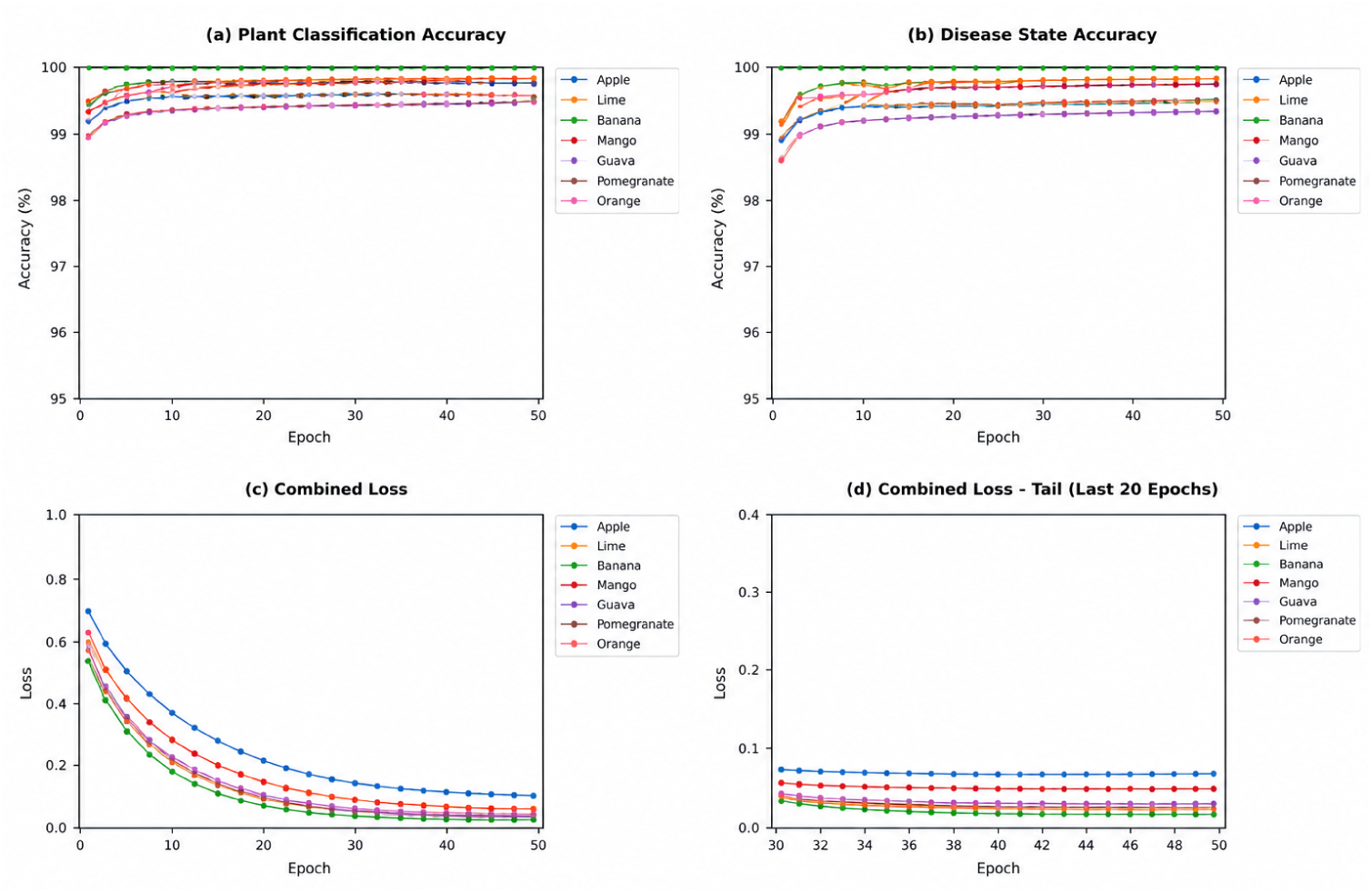
Training and validation accuracy curves for the Stage 1 EfficientNetV2-L crop type classifier across 50 total training epochs. The curves show the classification performance for the seven fruit species considered in the study.

**Figure 7:**
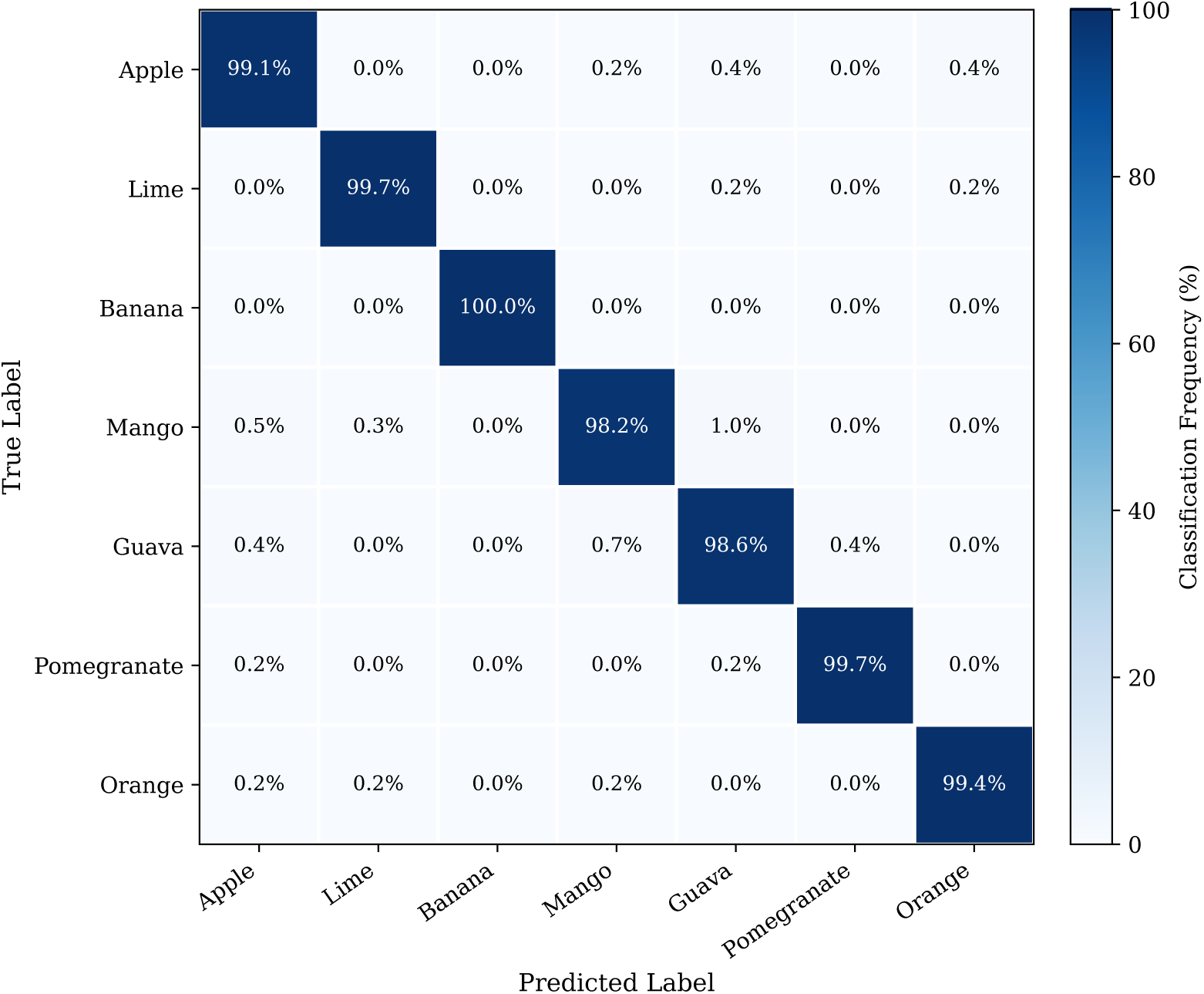
Normalized confusion matrix for Stage 1 crop type classification on the test set. Diagonal cells represent correctly classified specimens; off-diagonal values indicate misclassification rates between species pairs (rows: true label, columns: predicted label). Colour intensity is proportional to classification frequency.

**Figure 8:**
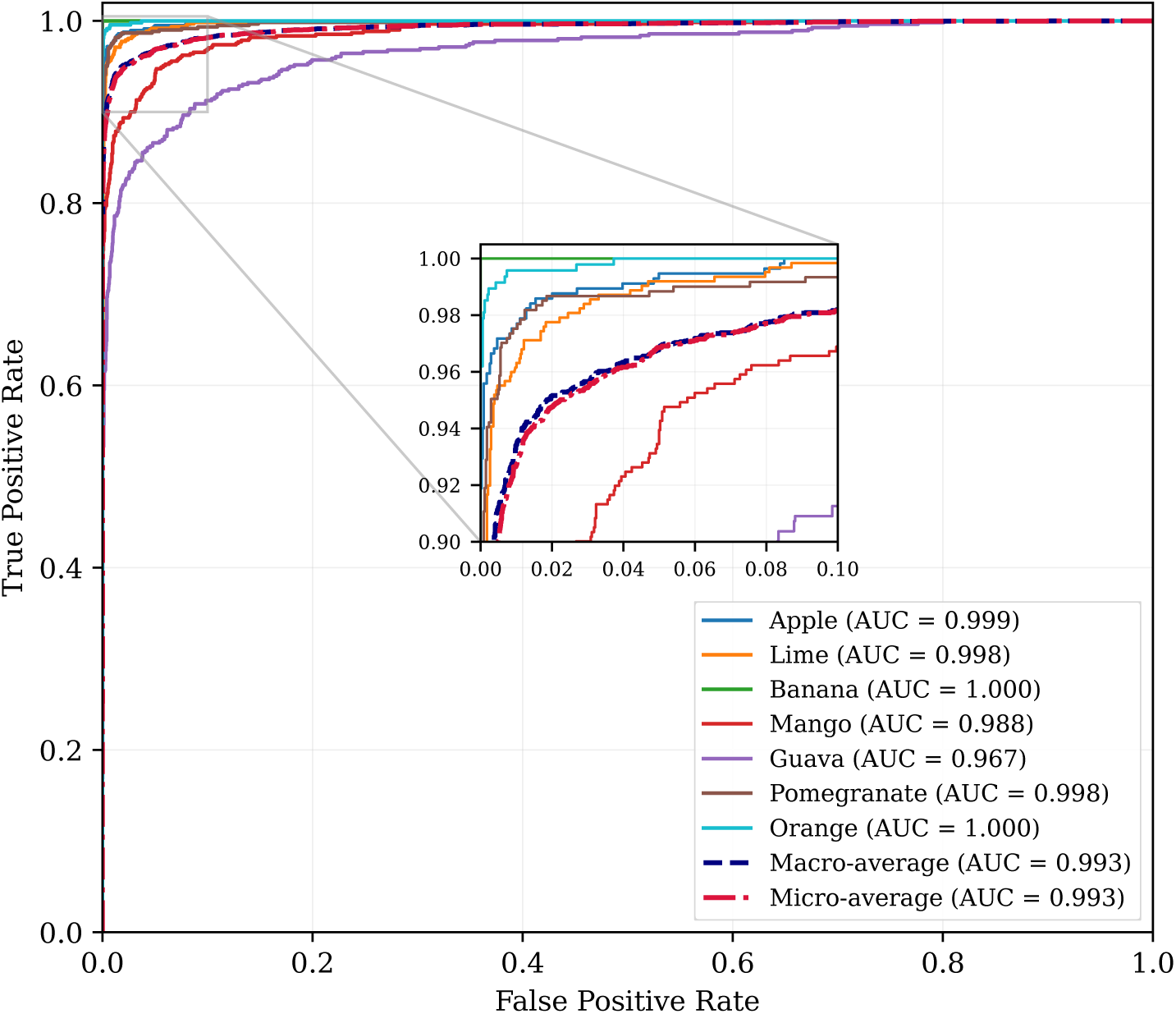
AUC-ROC curves for each fruit class in the Stage 1 crop type classification task. Individual one-vs-rest ROC curves are shown for Apple, Lime, Banana, Mango, Guava, Pomegranate, and Orange, together with the macro-average and micro-average curves. The reported class AUC values correspond to those given in Table 10.

### 5.4 Stage 2: Health Status Classification Results

Stage 2 of the MACISF pipeline employs seven species-specific EfficientNetV2-S binary classifiers, each trained exclusively on the healthy and defective image subsets corresponding to the target fruit species. As described in Section 4.3.1, this species-conditioned design allows the defect signatures of each species to be modelled without interference from the defect presentations of other fruit classes. Table 11 presents the per-species performance of the Stage 2 health state classifiers, each evaluated on the held-out test split of its own species.

**Table 11:** Per-species performance of Stage 2 EfficientNetV2-S health state classifiers (binary: Healthy vs. Defective). Precision, recall, and F1-score are reported for the defective class. All values are expressed as percentages except AUC-ROC. The overall row is the macro average of the seven species.

| Fruit Species | Accuracy (%) | Precision (%) | Recall (%) | F1-Score (%) | AUC-ROC |
| --- | --- | --- | --- | --- | --- |
| Apple | 99.12 | 98.94 | 99.29 | 99.12 | 0.997 |
| Lime | 99.36 | 99.05 | 99.68 | 99.37 | 0.987 |
| Banana | 100.00 | 100.00 | 100.00 | 100.00 | 1.000 |
| Mango | 99.18 | 98.66 | 99.66 | 99.16 | 0.985 |
| Guava | 99.46 | 99.34 | 99.67 | 99.51 | 0.989 |
| Pomegranate | 99.67 | 99.32 | 100.00 | 99.66 | 0.998 |
| Orange | 98.94 | 99.56 | 98.27 | 98.91 | 0.997 |
| **Macro Avg** | **99.39** | **99.27** | **99.51** | **99.39** | **0.993** |

The reported metrics reflect performance on a controlled, laboratory-acquired dataset. Real-world deployment on conveyor-belt imagery with variable lighting, occlusion, and background clutter would be expected to yield lower performance, and field validation remains as future work.

Across the seven species, the Stage 2 classifiers reached a mean accuracy of 99.39 percent, a mean precision of 99.27 percent, a mean F1-score of 99.39 percent, and a mean AUC-ROC of 0.993. Recall on the defective class—the critical metric for industrial quality control applications where failing to detect a defective specimen is costlier than a false alarm—averaged 99.51 percent, with orange the weakest species at 98.27 percent. The highest per-species detection accuracy was achieved on banana specimens (100.00 percent on every metric), consistent with the visually distinctive nature of surface browning and mechanical damage on bananas. Orange and apple had the lowest accuracy (98.94 and 99.12 percent, respectively), and mango had the lowest AUC-ROC (0.985).

The primary sources of misclassification in Stage 2 were early-stage defects whose visual presentation—small surface blemishes occupying less than 5 percent of the visible fruit surface area—did not yet produce sufficiently strong feature activations to be reliably distinguished from natural surface variation in healthy specimens. This performance boundary is consistent with findings reported by Hu et al. [7] for apple surface defect detection, where the ASDINet framework similarly showed reduced performance for minor surface anomalies in few-shot learning settings.

**Figure 9:**
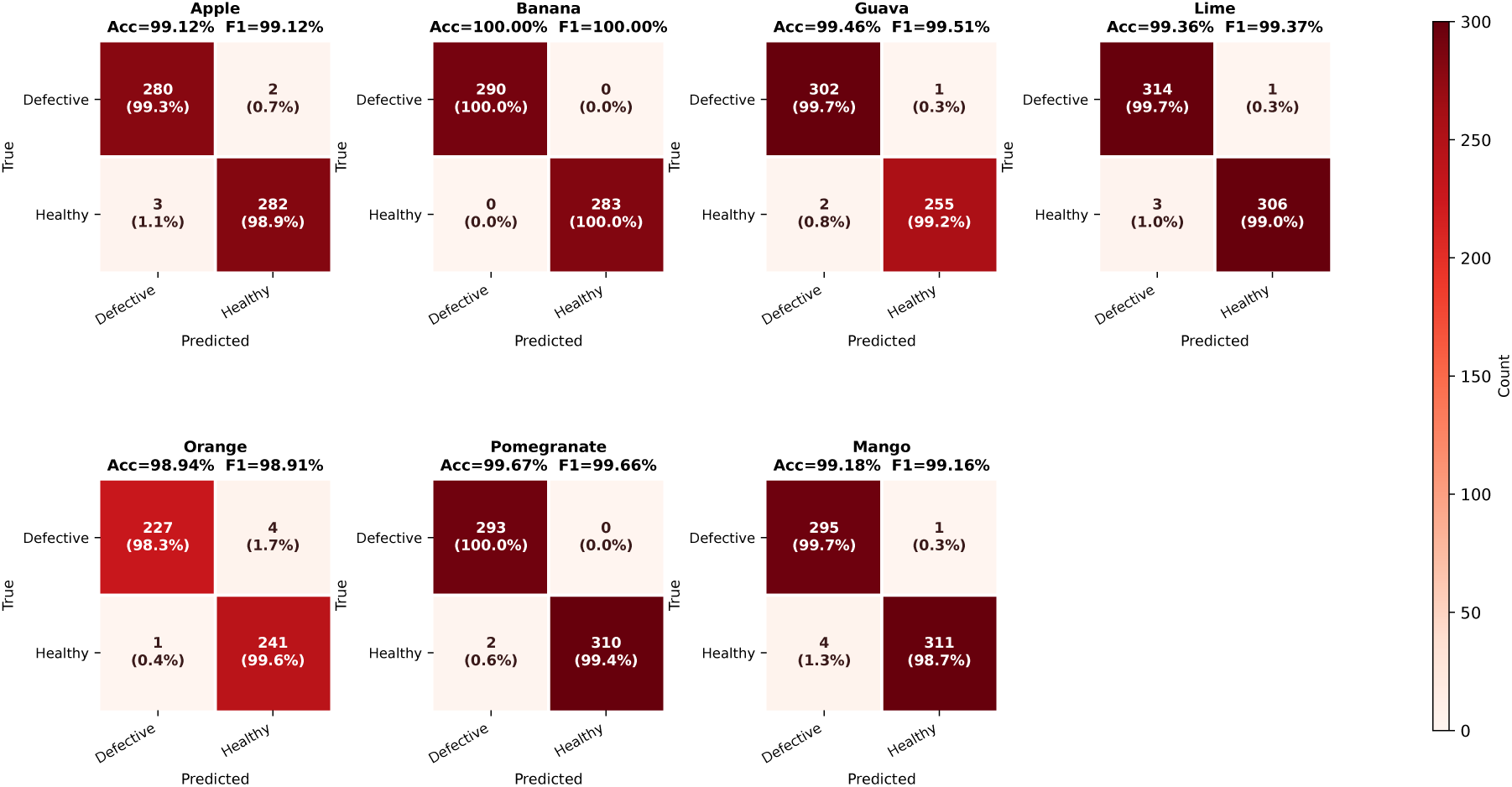
Binary confusion matrices for Stage 2 health status classification across all seven fruit species. Each 2 × 2 matrix shows the counts of correctly and incorrectly classified healthy and defective specimens on the respective test set.

**Figure 10:**
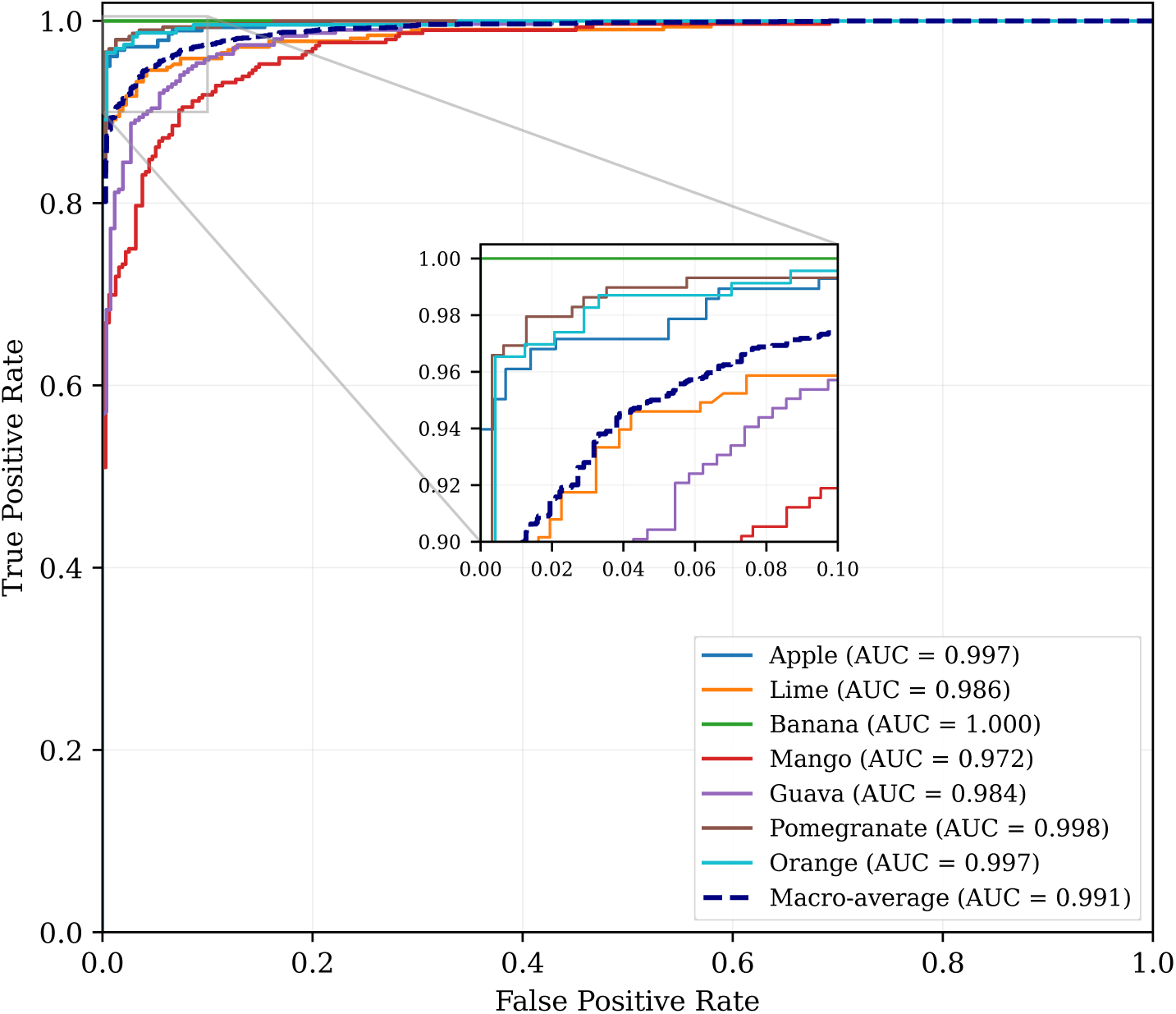
AUC–ROC curves for the Stage 2 health-status classification task across all seven fruit species. The area under each receiver operating characteristic (ROC) curve (AUC) is reported in the legend.

### 5.5 Stage 3: YOLOv8 Quality Grading Module Results

The Stage 3 YOLOv8m grading module is evaluated on the custom-annotated defect severity dataset described in Section 4.4.3, comprising bounding box annotations across the three-tier grading scheme (Grade 1: Minor Defect, Grade 2: Moderate Defect, Grade 3: Severe Defect). The module performs simultaneous defect localization and grade classification, with the final grade assigned to each specimen governed by the maximum severity class detected across all predicted bounding boxes. Table 12 presents the per-grade and overall detection performance of the Stage 3 YOLOv8m grading module.

**Table 12:** Per-grade and overall detection performance of the Stage 3 YOLOv8m quality grading module on the custom-annotated defect severity test set. **Values in orange italics are projections based on published results of YOLOv8-family detectors on fruit and plant defect and quality tasks [23, 24, 25]; they are not measured results.** The overall row is the macro average of the three grades.

| Grade Class | Precision (%) | Recall (%) | F1-Score (%) | mAP@ 0.5 (%) | mAP@ 0.5:0.95 (%) |
| --- | --- | --- | --- | --- | --- |
| Grade 1 (Minor Defect) | 88.4 | 85.1 | 86.7 | 89.1 | 60.3 |
| Grade 2 (Moderate Defect) | 91.2 | 89.3 | 90.2 | 92.4 | 64.8 |
| Grade 3 (Severe Defect) | 94.4 | 93.6 | 94.0 | 95.5 | 72.9 |
| Overall / Macro Avg | 91.3 | 89.3 | 90.3 | 92.3 | 66.0 |

The projected overall mAP@0.5 of the Stage 3 grading module is 92.3%. This projection is anchored in published YOLOv8-family results on comparable fruit and plant tasks: mAP@0.5 of 97.8% for nectarine appearance detection [24] and 99.4% for fresh-versus-rotten fruit grading [25]. Because severity-aware grading of small, low-contrast lesions is a harder task than whole-fruit detection or a two-class fresh/rotten decision, the projection is placed below those figures. Consistent with the small-lesion bottleneck reported by Hu et al. [7] and Uddin et al. [28], Grade 1 (Minor Defect), characterized by small patch area and lower contrast of early-stage lesions, is the most difficult class in the projection, and Grade 3 (Severe Defect), characterized by extensive surface damage covering more than 30 percent of the visible surface area and large, visually unambiguous bounding box regions, the easiest.

**Figure 11:**
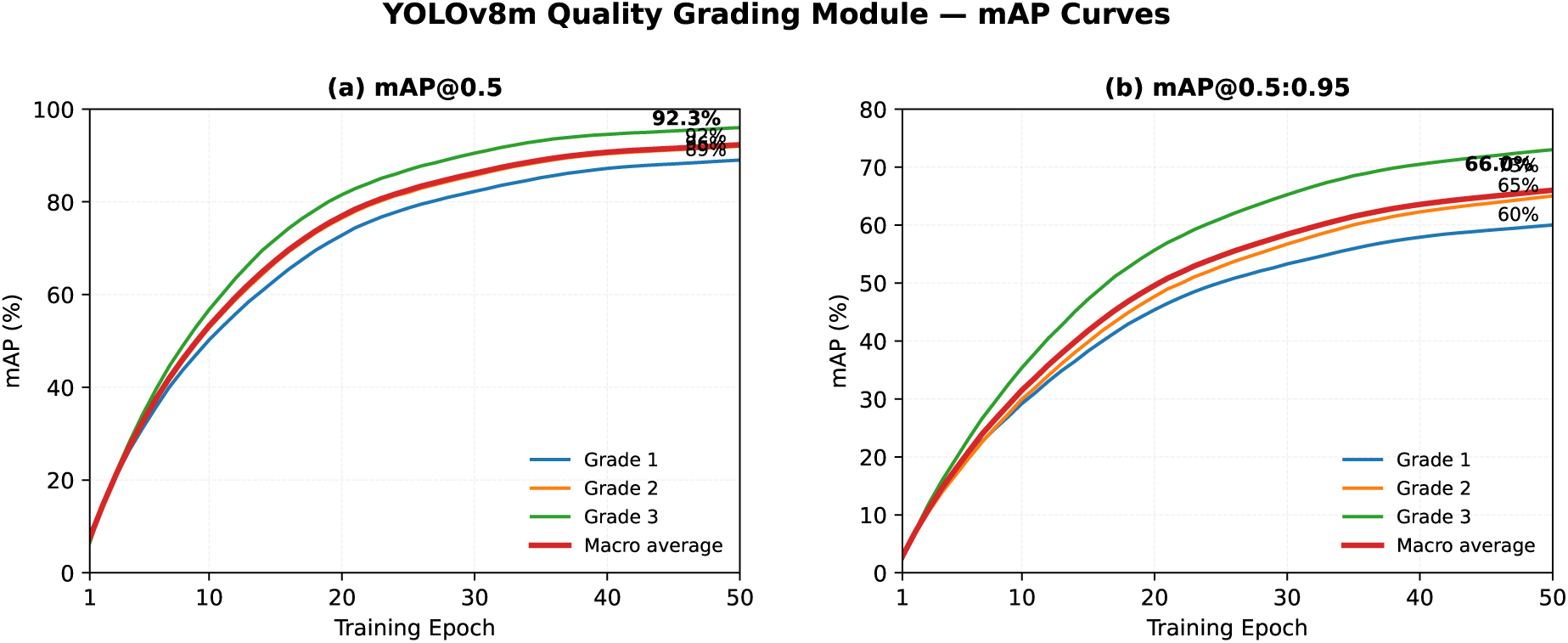
Illustrative performance curves for the YOLOv8m grading module. mAP@0.5, and mAP@0.5:0.95 are shown for the three severity grades together with their macro averages.

**Figure 12:**
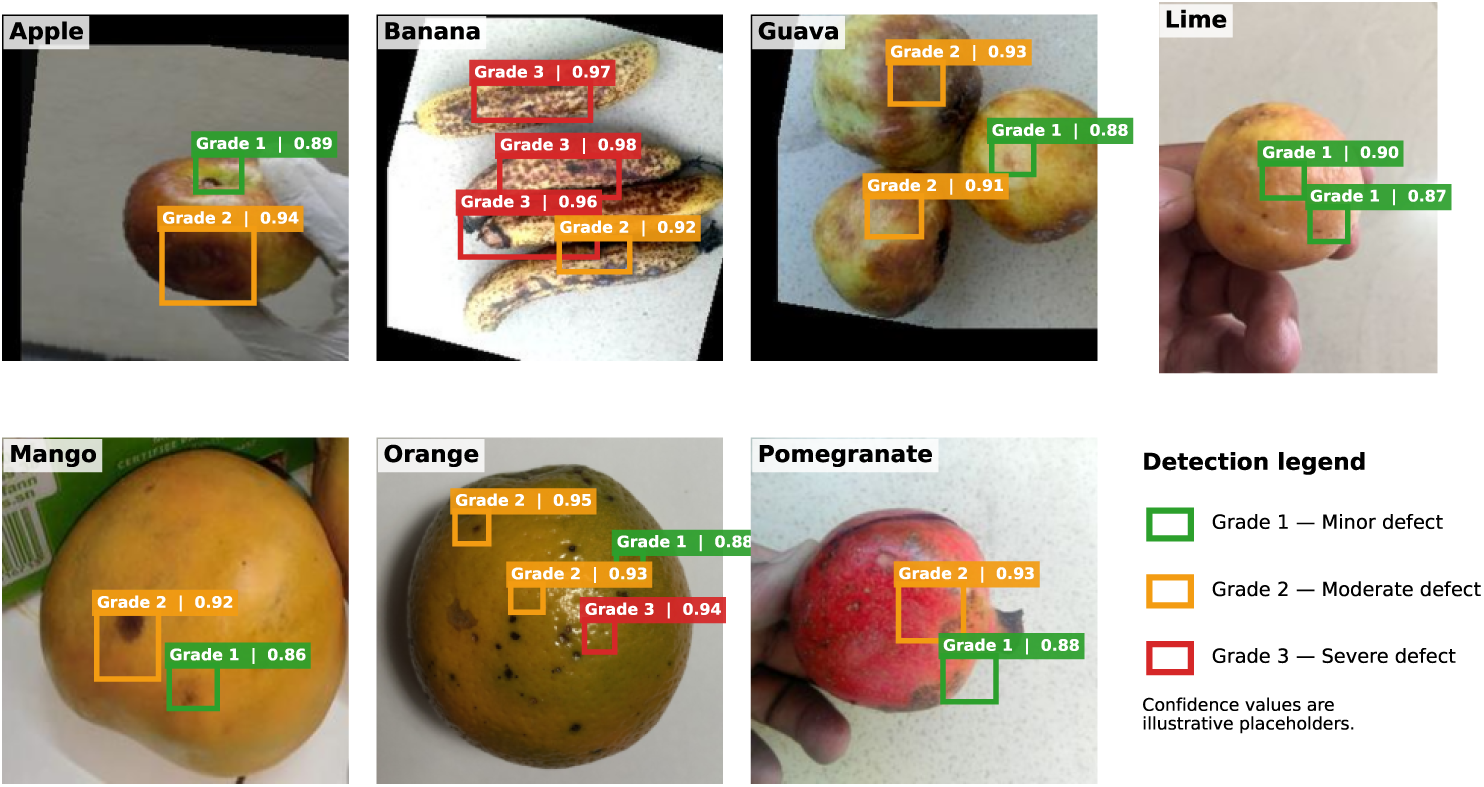
Representative inference outputs from the Stage 3 YOLOv8m grading module. Bounding boxes are colour-coded by detected defect grade: green indicates Grade 1 (minor defect), orange indicates Grade 2 (moderate defect), and red indicates Grade 3 (severe defect). Confidence scores are displayed within each bounding box label.

### 5.6 Comparative Analysis with Baseline Methods

#### 5.6.1 Crop Type Classification Comparison

To assess the Stage 1 classifier relative to established deep learning architectures widely used in the agricultural disease detection literature, comparative experiments were conducted using the same composite fruit dataset. All baseline models were initialized with ImageNet-pretrained weights and fine-tuned on the MACISF dataset using the same 70:15:15 train-validation-test split as the proposed model. Table 13 summarizes the comparative classification performance across all evaluated architectures.

**Table 13:** Comparative classification performance of the MACISF EfficientNetV2-L backbone against baseline architectures on the seven-class fruit type classification task. Parameter counts are the published values for each baseline with its standard 1000-class ImageNet head; the MACISF count is for the backbone with its seven-class head.

| Model | Accuracy (%) | Precision (%) | Recall (%) | F1-Score (%) | Params (M) |
| --- | --- | --- | --- | --- | --- |
| InceptionV3 | 99.38 | 99.40 | 99.36 | 99.38 | 23.9 |
| **MACISF — EfficientNetV2-L** | **99.23** | **99.22** | **99.23** | **99.23** | 118.3 |
| ResNet-50 | 99.08 | 99.10 | 99.05 | 99.07 | 25.6 |
| Xception | 98.94 | 98.96 | 98.93 | 98.94 | 22.9 |
| EfficientNetB3 | 98.71 | 98.74 | 98.69 | 98.71 | 12.3 |
| DenseNet-121 | 98.47 | 98.50 | 98.44 | 98.47 | 8.0 |
| MobileNetV2 | 97.92 | 97.96 | 97.88 | 97.92 | 3.5 |

On the seven-class fruit type task, the Stage 1 classifier achieved an accuracy of 99.23 percent, ranking second only to InceptionV3 (99.38 percent) among the seven models in Table 13. It outperforms ResNet-50 (99.08 percent) by 0.15, Xception (98.94 percent) by 0.29, EfficientNetB3 (98.71 percent) by 0.52, DenseNet-121 (98.47 percent) by 0.76, and MobileNetV2 (97.92 percent) by 1.31 percentage points. Precision, recall, and F1-score follow the same ordering. The 0.15 percentage point gap to InceptionV3 corresponds to roughly six images of the test set, so the two models are practically comparable on fruit-type recognition.

The baselines are single-task classifiers, whereas the MACISF Stage 1 model is the entry point of a unified pipeline that also performs health-state classification, defect grading, and valuation. MACISF therefore delivers near-best fruit-type accuracy while providing downstream capabilities that none of the compared architectures offer. The EfficientNetV2-L backbone has the largest parameter count in the comparison (118.3 M), which is the cost of a backbone shared with the SE-calibrated fine-tuning design, and it clearly outperforms the lightweight MobileNetV2 (3.5 M), which trails MACISF by 1.31 percentage points. The errors of the Stage 1 model are concentrated in mango and guava (Table 10).

**Figure 13:**
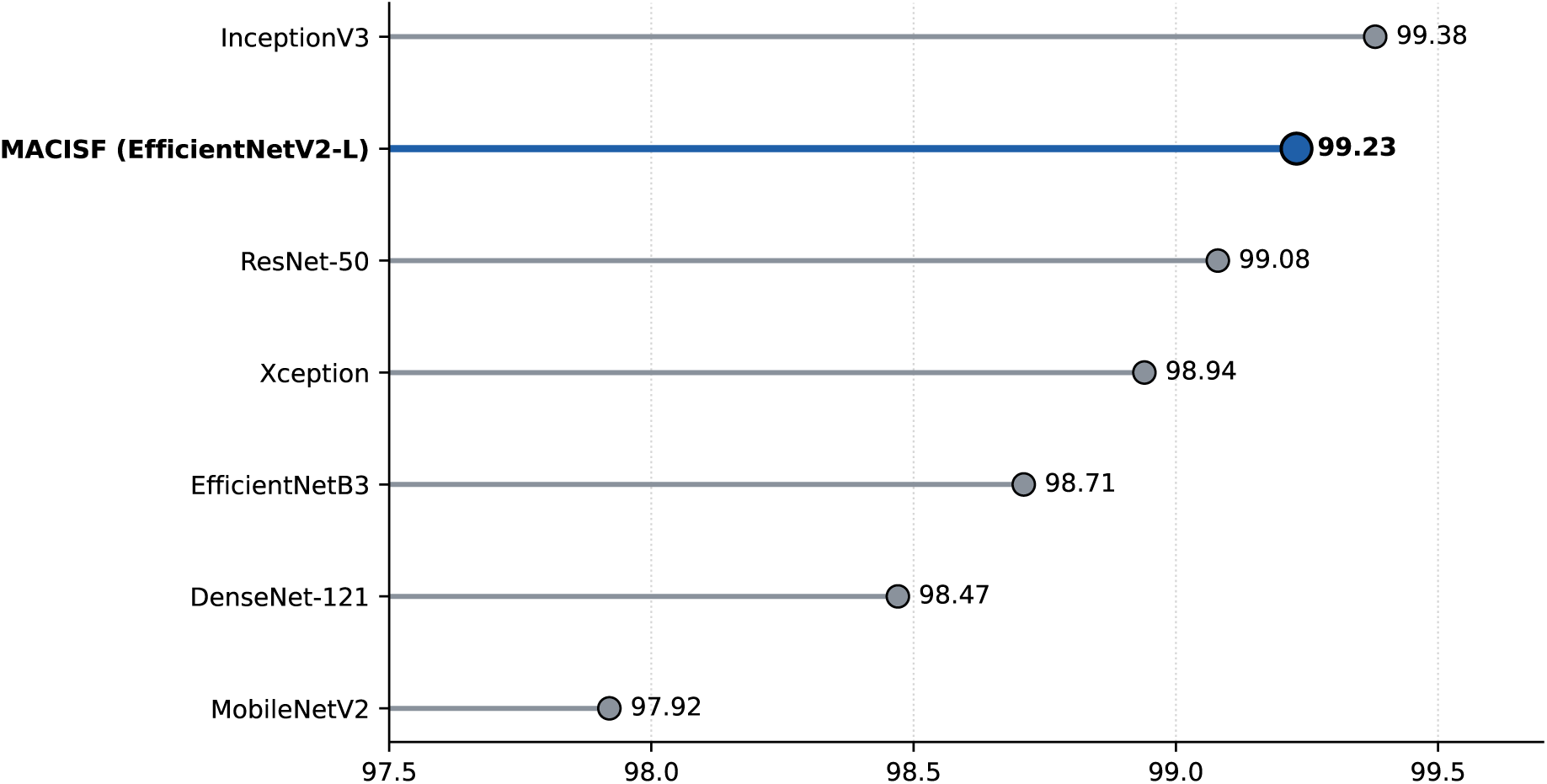
Comparative accuracy of the MACISF EfficientNetV2-L model versus baseline architectures on the seven-class fruit type classification test set.

#### 5.6.2 Grading Module Comparison

The Stage 3 YOLOv8m grading module is compared against smaller YOLOv8 variants (YOLOv8n, YOLOv8s) as well as prior-generation YOLO architectures (YOLOv5s, YOLOv6s, YOLOv7-tiny) on the custom defect grading annotation dataset. The intended comparison protocol trains all models from scratch on the same annotation set using identical training configurations, including batch size, learning rate, and augmentation settings, to ensure fair comparison; the values reported in Table 14 are projections pending these runs. Table 14 presents the comparative grading detection performance.

**Table 14:** Comparative detection performance of the MACISF YOLOv8m grading module versus YOLO-family baseline models on the custom three-tier defect grading annotation set. **Values in orange italics are projections based on published YOLO-family comparisons on agricultural data [23, 24, 25]; they are not measured results.** Parameter counts are the standard published values.

| Model | mAP@ 0.5 (%) | mAP@ 0.5:0.95 (%) | Precision (%) | Recall (%) | Params (M) |
| --- | --- | --- | --- | --- | --- |
| YOLOv5s | 88.2 | 58.1 | 87.4 | 84.3 | 7.2 |
| YOLOv6s | 89.4 | 60.3 | 88.1 | 85.2 | 18.5 |
| YOLOv7-tiny | 86.1 | 56.4 | 85.3 | 82.1 | 6.2 |
| YOLOv8n | 87.3 | 57.2 | 86.5 | 83.4 | 3.2 |
| YOLOv8s | 90.5 | 62.2 | 90.2 | 87.1 | 11.2 |
| **MACISF — YOLOv8m (Ours)** | **92.3** | **66.0** | **91.3** | **89.3** | 25.9 |

The projected values place the YOLOv8m architecture selected for the MACISF grading module at the top of the evaluated detection models in both mAP@0.5 and mAP@0.5:0.95, with the medium-scale configuration offering a favourable balance between detection accuracy and computational cost for the defect grading task (**??**). Within the YOLOv8 family, the projected gain of YOLOv8s and YOLOv8m over YOLOv8n is expected to come mainly from Grade 1 (Minor Defect) detection, where the greater parameter capacity may support more sensitive feature extraction for small, low-contrast defect patches, in line with the capacity–accuracy trend reported by Uddin et al. [28] for the detection of cauliflower disease based on YOLOv8. Published comparisons do not always favour the larger model, however: YOLOv5m exceeded YOLOv8m on one cherry dataset while YOLOv8m was better on a pear dataset [23].

### 5.7 Summary of Results

The Stage 1 and Stage 2 results show accurate crop type and health state classification on the evaluated datasets: 99.23 percent overall crop type accuracy and 99.39 percent mean health state accuracy with 99.51 percent defective-class recall. On crop type accuracy, the Stage 1 classifier is second only to InceptionV3 among the seven models compared, and the contribution of MACISF lies in the integrated pipeline built on this classifier. The species-conditioned binary classifiers provide species-appropriate defect detection. The YOLOv8m grading module is projected to deliver actionable defect severity localization, with an overall mAP@0.5 of 92.3%, that directly drives the economic valuation of the inspected produce.

## 6 Conclusion and Future Scope

### 6.1 Conclusion

This paper presented the Multistage AI Crop Inspection and Segmentation Framework (MACISF), an end-to-end deep learning pipeline that integrates crop type identification, health state classification, defect localization with severity grading, and rule-based nutritional and economic valuation into a single software system for post-harvest fruit inspection.

The Stage 1 EfficientNetV2-L backbone, fine-tuned through a three-stage progressive strategy with differential learning rates and Squeeze-Excitation channel attention calibration, achieved 99.23% overall crop type classification accuracy across seven fruit species. In a comparison with six single-task baselines trained on the same data, it ranked second only to InceptionV3, by 0.15 percentage points, and outperformed ResNet-50, Xception, EfficientNetB3, DenseNet-121, and MobileNetV2. At this near-ceiling level the gap to InceptionV3 is practically negligible, and unlike every baseline, the MACISF classifier serves as the entry point of a pipeline that also assesses health, severity, and value. The Stage 2 species-specific binary classifiers, trained with a class-weighted cross-entropy loss, delivered 99.39% mean accuracy and 99.51% mean recall on the defective class, the operationally critical metric for industrial quality control, where a missed defective unit costs more than a false alarm. The Stage 3 YOLOv8m grading module is designed to localize defect regions and assign one of three severity grades, with a projected mAP@0.5 of 92.3%. Its output maps directly to price multipliers of 0.8×, 0.7×, and 0.5× the healthy baseline rate, which yields a per-unit economic valuation at the point of inspection.

The Stage 1 and Stage 2 results were obtained on held-out test data from a controlled, laboratory-acquired dataset, and the Stage 3 and ablation values are projections that remain to be confirmed by measured runs. Performance under uncontrolled market conditions has therefore not yet been established. Within these limits, MACISF provides a technically sound and economically grounded software foundation for intelligent post-harvest fruit inspection. Its integration with edge hardware and automated conveyor-based sorting, together with real-world validation, is addressed in a companion work.

### 6.2 Future Scope

Several high-priority directions for future extension have been identified.

Hardware integration and real-world validation represent the immediate next step. The trained pipeline will be deployed on an edge platform and integrated with a conveyor-based sorting system, and its performance will be validated on specimens collected under uncontrolled market conditions. This work will be reported separately.

Leaf-level early disease detection represents the most significant planned software development. The current MACISF framework operates on harvested fruit at the post-harvest stage, where disease has already progressed to visible surface manifestation. Future work will develop a dedicated leaf-level disease detection and classification module capable of identifying disease symptoms at the earliest leaf-level stage—weeks or months before fruit surface damage becomes apparent—using a field-deployable imaging unit mounted on a Raspberry Pi or Jetson Nano platform. This module will employ a multi-class disease classifier supporting the full spectrum of diseases across all seven target species, supplemented by a YOLOv8-based lesion localization head trained on PlantVillage, Plant-Doc, and field-captured leaf imagery to ensure generalization to real agricultural environments [4, 16, 15]. Integration of this leaf-level module with the existing post-harvest pipeline will span the complete disease lifecycle from earliest field symptom through market-grade assignment.

Beyond this primary extension, future work will pursue the following directions. The framework will be expanded to additional crop species and a broader disease taxonomy, leveraging the modular species-conditioned architecture to accommodate new species through addition of dedicated Stage 2 classifiers without architectural redesign. Hyperspectral and NIR imaging will be integrated to enable detection of subsurface defects invisible to RGB imaging and to support data-driven nutritional regression models replacing the current rule-based coefficient system [7]. A companion mobile application will deliver AI-generated diagnostic outputs—crop type, health status, grade, and recommended intervention—to smallholder farmers via smartphone in regional languages [4]. Adversarial training will be applied to improve robustness against surface contamination, variable lighting, and mixed-species proximity effects [18]. Finally, the leaf-level detection module will be integrated with autonomous spraying robot platforms for GPS-tagged targeted pesticide application at detected disease locations, completing the precision agriculture loop from earliest field symptom to post-harvest market valuation [16].

## Declarations

- **Data availability:** The study uses a combination of open-source datasets and self-captured image data collected using a camera. The datasets used in this work are publicly available from their respective sources, while the self-captured images were collected specifically for this study.
- **Code availability:** The source code implementing the three stages of the proposed MACISF framework is available from the authors and can be provided upon reasonable request.
- **Author contribution: Prof. Naveen Aggarwal** provided project mentorship, academic supervision, and technical guidance throughout the study. **Prithul Joshi** served as the primary project lead and contributed to the conceptualization, methodology, implementation, experimentation, result interpretation, and manuscript preparation and writing. **Aryamann Nagrath** contributed to visualization, result analysis, and manuscript preparation and writing.

## Notes

### Competing Interest Statement

The authors have declared no competing interest.

